# Spatial profiling of CAR protein organization reveals in vivo remodeling during CAR-T therapy

**DOI:** 10.64898/2026.04.20.719384

**Authors:** Yukie Kashima, Kenichi Makishima, Hanna van Ooijen, Lovisa Franzén, Stefan Petkov, Hidekazu Nishikii, Junko Zenkoh, Ayako Suzuki, Annika Branting, Mamiko Sakata-Yanagimoto, Yutaka Suzuki

**Author notes:** Co-corresponding authors, Yutaka Suzuki, Life Science and Data Science Center, Graduate School of Frontier Sciences, The University of Tokyo, Chiba, Japan, 5-1-5, Kashiwanoha, Kashiwa, Chiba 277-8562, Japan. Contribute equal to this study (Yukie Kashima, Kenichi Makishima, Hanna van Ooijen).

## Abstract

Chimeric antigen receptor (CAR) T cell therapy utilizes genetically engineered patient-derived T cells to target cancer cells. Despite its clinical successes in multiple cancer types, the underlying molecular mechanisms by which molecules on CAR-T cells and surrounding cells interact with other proteins and collectively determine treatment efficacy remain elusive. Most previous studies have relied on transcriptome profiling, which does not fully reflect protein-level organization and interactions. In this study, we developed an antibody-oligonucleotide conjugate targeting the FMC63 region of CAR and integrated it into molecular pixelation (MPX). This approach enabled profiling of the dynamics of CAR molecules on cell surfaces as well as their colocalization with other proteins at the single-cell level. By applying MPX to longitudinal samples from three patients undergoing CAR-T cell therapy, we characterized the dynamic changes in CAR-associated protein organization in both pre-infusion CAR products and post-infusion peripheral blood. While CAR protein abundance and polarization showed limited variation across clinical courses, remodeling of a CAR-centered co-localization network was observed over time, including different retentions of specific molecular associations between patients with different clinical outcomes. Although derived from a limited cohort, our study identifies insights from this methodological framework beyond those gained by conventional omics analyses and offers results of a systematic investigation to predict and enhance CAR therapeutic outcomes.

**Key points:**

- Molecular pixelation was applied for chimeric antigen receptor (CAR) profiling at single-molecule and single-cell resolutions.
- Protein and transcriptome analyses of the CAR molecule showed dynamic remodeling during CAR-T therapy in patients with non-Hodgkin lymphoma.

## Introduction

Chimeric antigen receptor (CAR) T cell therapy has emerged as a potential immunotherapeutic strategy for cancer. In this approach, autologous T cells are engineered to express CAR, including an endodomain, an anchoring transmembrane domain, and an ectodomain^1^. CAR-T cell therapies targeting CD19 have been successfully applied in patients with relapsed or refractory (R/R) B cell hematological malignancies such as B cell acute lymphocytic leukemia and B cell non-Hodgkin’s lymphoma^2^, demonstrating remarkable clinical outcomes. Several CD19-targeting CAR-T products, including tisagenlecleucel (tisa-cel: Kymriah), axicabtagene ciloleucel (axi-cel: Yescarta), and lisocabtagene maraleucel (liso-cel: Breyanzi), have been developed and are widely used for the treatment of B cell malignancies. Tisa-cel was the first CD19-targeting CAR-T cell therapy^3^ approved by the U.S. Food and Drug Administration, and it is currently approved for R/R B cell acute lymphoblastic leukemia, R/R diffuse large B cell lymphoma, and R/R follicular lymphoma^4^.

Despite its overall success, CAR-T cell therapy faces several critical challenges. A substantial proportion of patients develop cytokine release syndrome (CRS), which can necessitate discontinuation of treatment. Moreover, long-term responses are not achieved in all patients, and relapse remains common. Relapse has been linked to antigen loss (e.g., CD19 loss in tumor cells), the complexity of the tumor microenvironment, molecular landscape, and CAR-T heterogeneity^5–7^. Early clinical trials in solid tumors demonstrated limited efficacy compared to hematologic malignancies^8^. Although these factors have been extensively investigated, the mechanisms underlying response and relapse remain incompletely understood.

Recent studies highlighted marked heterogeneity within CAR-T populations^9,10^. In some cases, patient-to-patient variability in CAR-T cell abundance reflects differences in pre-manufacturing T cell states as well as dynamic changes during CAR-T cell production. Longitudinal analyses following CAR-T cell infusion have demonstrated distinct population dynamics in peripheral blood, with expansion and persistence patterns associated with clinical response^11^. However, the molecular features distinguishing effective from ineffective CAR-T states in pateints remain incompletely characterized. Moreover, transcript expression does not reflect protein expression or spatial organization at the cell surface. CAR-specific features—including receptor abundance, spatial distribution, and protein interactions—remain largely unexplored at single-cell level. Although several studies have used CAR-specific antibodies to assess protein expression, they primarily capture bulk- or population-level changes, providing limited insight into single-cell-resolved protein abundance, spatial distribution, and cell interactions.

To address this gap, we applied molecular pixelation (MPX), a recently developed technology^12^ that enables high-resolution profiling of cell surface proteins at single-cell resolution. Here, we provide a detailed characterization of CAR-T cell surface protein dynamics using a CAR-specific antibody-oligonucleotide conjugate (AOC) targeting the FMC63 single-chain variable fragment (scFv) region widely used in CD19-targeting CAR-T^13^ therapies. We quantified surface protein abundance and spatial organization at single-cell resolution and examined longitudinal changes across patients with different clinical courses. Integrating these protein-level findings with single-cell RNA-seq, we interpret molecular events associated with CAR-T treatment.

## Results and Discussion

### MPX system profiling CAR-T molecules via FMC63

Current MPX AOCs include 76 key cell-surface proteins on major immune cells: T, B, NK, and monocytes. To enable an analysis of CAR-T cell-specific protein features, we first developed a novel AOC targeting FMC63 scFv that serves as the extracellular antigen-binding domain of Tisa-cel^14,15^ (Figure 1A and 1B).

**Figure 1.**
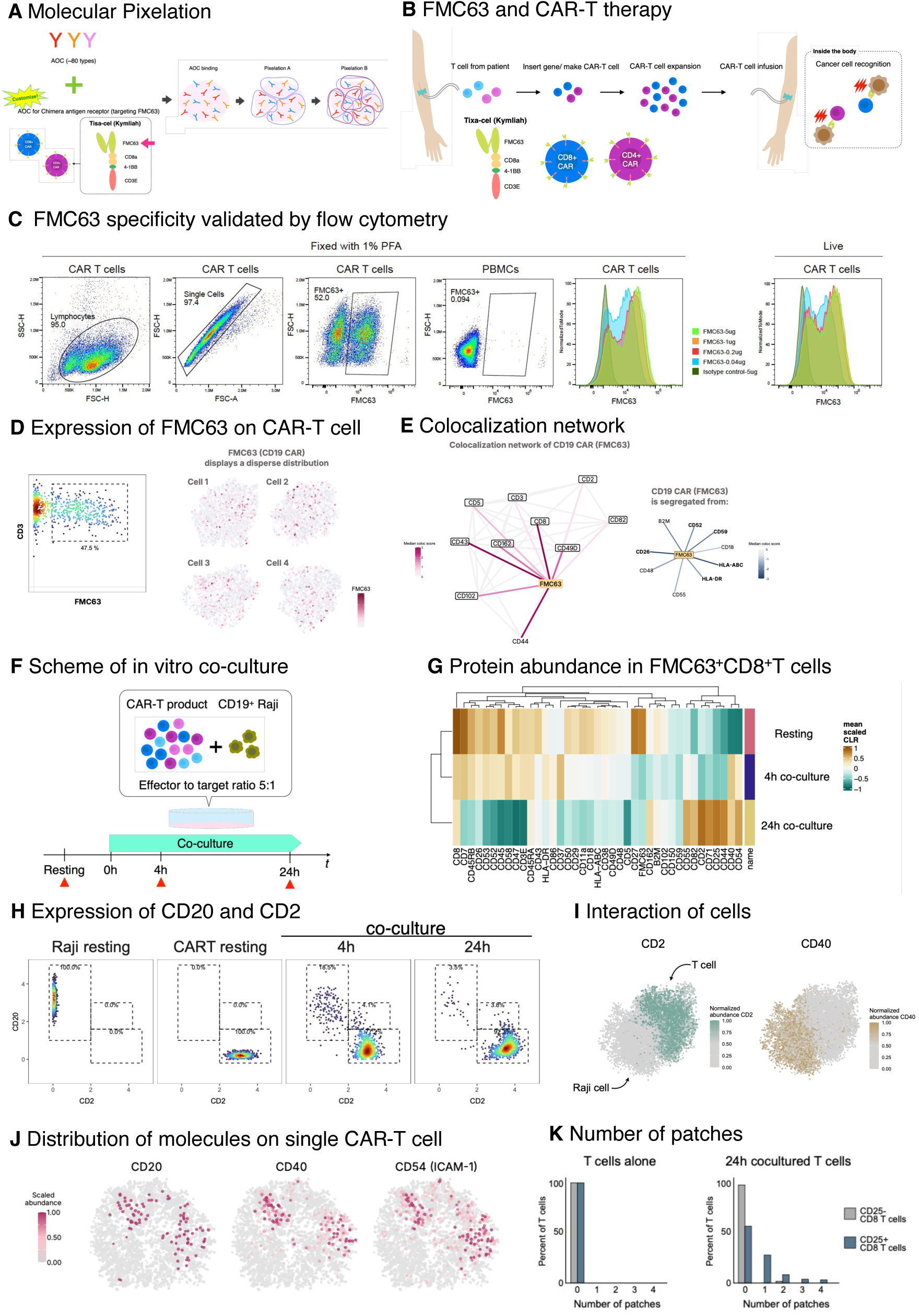
In vitro validation of FMC63 as a CAR detection target. **a,** Schematic overview of molecular pixelation (MPX) analysis workflow. **b**, Schematic illustration of CAR-T cell therapy. **c,** Validation by flow cytometry using 1% PFA fixed and live samples. The gating strategy used to identify CAR^+^ T cells and percentage of gated positive cells within the indicated gate are shown in each plot. FMC63-based CAR^+^ T cells were used as a positive control, and PBMCs from a CAR-therapy-naïve donor served as negative control. The FMC63 antibody concentration was 1 µg/mL (left). Representative histograms showing dose-dependent detection of CAR-positive T cells using FMC63 antibody at 5 µg/mL, 1 µg/mL, 0.2 µg/mL, and 0.04 µg/mL in fixed (left) and live (right) samples. **d**, Expression of FMC63 in single cells. Representative image of the gating used to identify FMC63^+^CD3^+^ cells in CD19-targeting CAR-T cells (left) and the distribution of FMC63 molecules in each cell. **e**, Co-localization network of CD19-targeting CAR-T cells. **f,** Schematic overview of the in vitro co-culture assay. CD19-targeting CAR T cells were co-cultured with CD19^+^ Raji cells at an effector-to-target ratio of 5:1. Samples were collected at rest (CAR-T product) and at 4 and 24 h of co-culture. **g**, Heatmap showing protein abundance in CD8^+^ CAR^+^ T cells. The selected markers are indicated. **h**, Representative gating plot of CD2 (x-axis) and CD20 (y-axis) in Raji cells and resting CAR-T cells at 4 and 24 h after co-culture (from left to right). The percentage of cells within each gate is shown. Identical thresholds were applied to all samples. **i**, Visualization of Raji and T cells from co-cultured samples, indicated by CD2 (a vital surface marker of T cells) and CD40 (a canonical marker for Raji cells). Two adjacent cells undergoing trogocytosis. **j,** Single-cell visualization of molecular distribution. Representative plots showing the spatial distribution of CD20, CD40, and CD54 on cell surfaces. Each panel corresponds to the same cell and is colored according to the expression levels of the indicated molecules. **k,** Histogram showing the distribution of the number of patches at rest (left) and at 24 h of co-culture (right).

The suitability of targeting FMC63 was first assessed by flow cytometry using commercial CD19-targeting CAR-T products. FMC63 detected 51.2 % of FMC63^+^ CAR^+^ cells after fixation with 1 % paraformaldehyde (PFA) at an antibody concentration of 1 µg/mL (Figure 1C). We next evaluated FMC63 staining across a range of antibody concentrations (5, 1, 0.2, 0.04 µg/mL) in both 1 % PFA fixed and live cells (Figure 1C). In contrast, PBMCs from a CAR-T therapy-naïve donor used as negative control were 99.9 % negative for FMC63 antibody staining (Figure 1C, PBMCs panel). This analysis confirmed that FMC63 specifically detected CAR-expressing T cells.

Next, performance and specificity of the MPX system incorporating FMC63 AOC were validated. The MPX assay was performed using FMC63 AOC and AOCs for the other 76 proteins. An abundance-based assay identified 47.5 % of cells as FMC63^+^CD3E^+^ T cells within a commercial CD19-targeting CAR-T product (Figure 1D), consistent with previous studies that showed CAR transduction efficacies of approximately 60–65 % in healthy and ∼30 % in patient-derived T cells^16^.

Under unstimulated conditions, FMC63 exhibited an almost uniform distribution on cell surfaces (Figure 1D). Co-localization network analysis revealed that FMC63 co-localized tightly with CD8 and CD43. In contrast, FMC63 was segregated from HLA proteins and beta 2 microglobulin (B2M) protein (Figure 1E). Collectively, these results indicate that active CAR was expressed on T cell surfaces.

To further evaluate performance of the FMC63 AOC, we performed an in vitro co-culture assay using CAR-T and CD19^+^ Raji cells. Cells were collected at 0, 4, and 24 h after co-culture and subjected to MPX assay (Figure 1F). Protein abundance of FMC63 on CAR-T cells, as well as that of CD8 and CD43, decreased during interactions with their target CD19 proteins (Figure 1G), indicating that CAR-T cells began losing their cytotoxic activity during intensive interactions with their target Raji cells.

Interestingly, CD20 abundance was significantly increased on the surfaces of CAR-T cells, even though this protein is a cell surface marker of Raji cells (Figure 1H). Further examination revealed that some CAR-T cells were physically interacting with Raji cells through a phenomenon known as “trogocytosis.” Cells undergoing trogocytosis were detected as doublet cells during gating in the abundance analysis (Figure 1H and 1I) and peaked at 4 h of co-culture (Figure 1H). Figure 1J shows representative CAR-T cell-exhibiting patches, defined as localized clusters of CD20, CD40, and CD54 (ICAM-1) signal intensity on CAR-T cell surfaces. These patches likely represent traces of prior interactions with Raji cells. Consistently, the number of FMC63^+^ CD8^+^ T cells with such patches increased after 24 h of co-culture. Interestingly, this patch was observed more frequently in CD25^+^ CD8^+^ T cells, a marker of activation (Figure 1K).

### Application of the MPX assay with FMC63 to clinical samples

To assess the performance of the FMC63 AOC in clinical samples, we performed validation analyses using multiplexed immunocytochemistry.

We recruited three patients with non-Hodgkin lymphoma who had received Tisa-cel therapy at a single institution (Figure 2A). All patients (donor A, B, and C) developed cytokine release syndrome (CRS) at days 0, 2, and 1 after infusion, respectively, and were treated with four, two, and one administration of tocilizumab, with one patient (donor C) additionally also administered 9.9 mg dexamethasone. One patient (donor C) experienced disease relapse 73 days after infusion, whereas the remaining two donors (donors A and B) maintained remission. Detailed clinical characteristics are provided in Supplementary Table S1.

**Figure 2.**
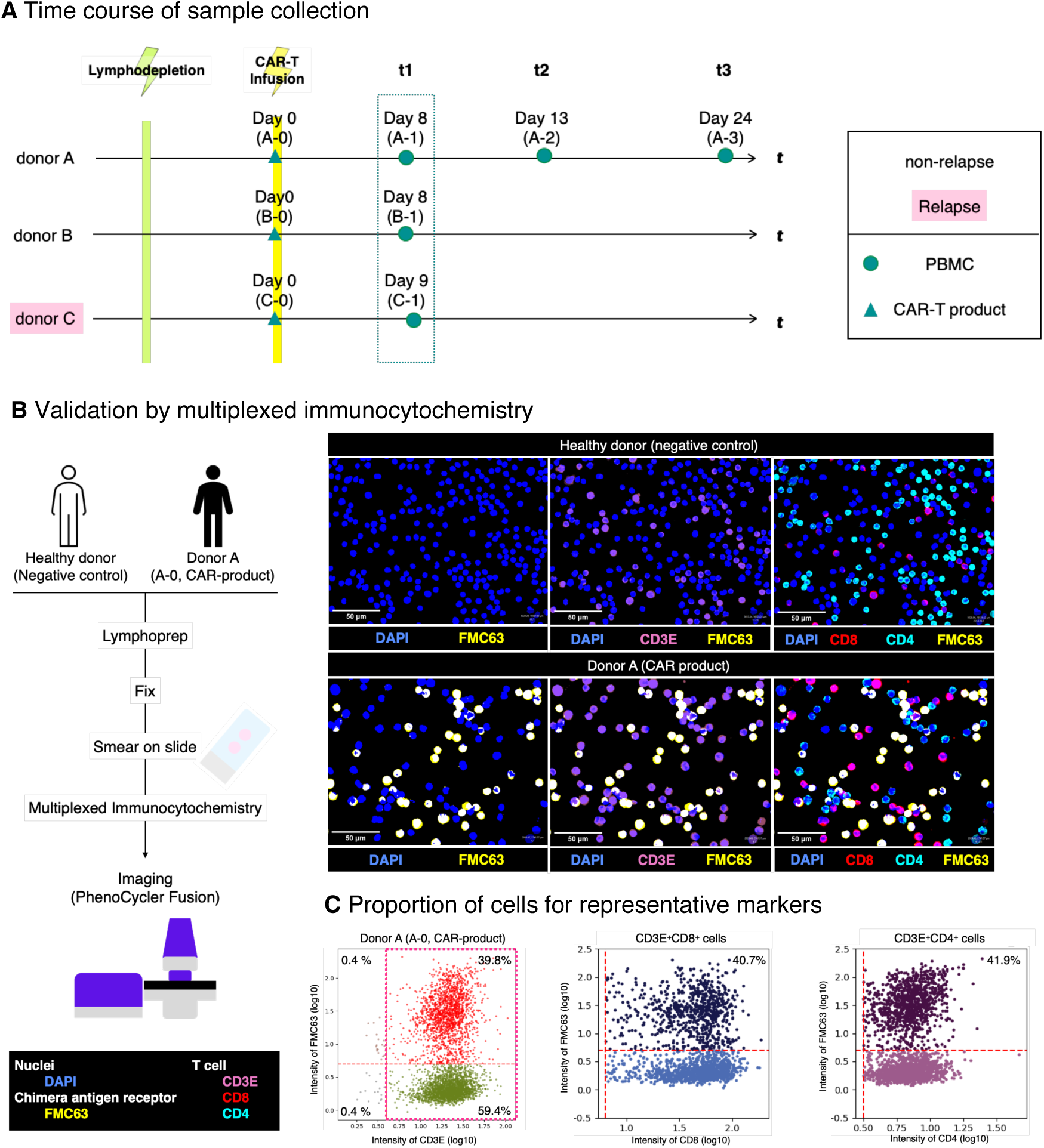
Validation of FMC63 in clinical samples. **a,** Timeline of sample collection from the three donors included in this study. **b**, Representative multiplexed immunocytochemical images stained for DAPI, CD3E, CD8, CD4, and FMC63. Merged images of cells from a healthy donor (FMC63 negative control) and donor A (FMC63 positive control) are shown. Scale bar is 50 µm. **c,** Validation using PhenoCycler -Fusion focusing on FMC63, CD3E, CD8, and CD4 proteins. The gating plot shows the proportions of cells in the indicated areas. The sample shown is from donor A (A-0, CAR-product).

CAR-T cell infusion products and PBMCs were analyzed. Donor A provided four samples, including the original CAR-T cells (A-0) and PBMCs collected on days 8 (A-1), 13 (A-2), and 24 (A-3) post-infusion. Donors B and C provided CAR-T products (B-0 and C-0), with PBMCs collected on days 8 (B-1) and 9 (C-1), respectively (Figure 2A and Supplementary Table S1). These time points were selected to capture the early post-infusion phase, during which CAR-T cell expansion typically peaks between 7 and 14 days^17,18^.

To further validate the FMC63 AOC in clinical samples, we performed multiplexed immunocytochemistry using a PhenoCycler-Fusion system with a customized antibody panel that included FMC63 and another 18 typical PBMC markers (Supplementary Table S2 and S3). The CAR-T cell product from donor A served as a positive control, whereas PBMCs from a healthy donor were used as a negative control. Clear FMC63 staining was observed exclusively in the donor A sample (Figure 2B), with 39.8 % of FMC63^+^CD3E^+^ cells, 40.7 % of FMC63^+^CD8^+^T cells (among CD3E^+^CD8^+^ T cells), and 41.9 % of FMC63^+^CD4^+^ T cells (among CD3E^+^CD4^+^ T cells), demonstrating FMC63 AOC specificity and sensitivity in clinical samples (Figure 2C). Channel information is shown in Supplementary Figure S1 and S2.

### Cellular composition and protein abundance profiles of pre-infusion CAR-T products

We performed an MPX analysis of clinical samples from patients who received CAR-T cell therapy using a panel of 76 AOCs supplemented with FMC63 AOC. In parallel, single-cell RNA-sequencing (scRNA-seq) was performed to assess mRNA expression profiles (Figure 3A). Across ten samples, 6,380 cells were analyzed using the MPX assay and 44,372 cells using the scRNA seq. Cells annotated as major immune cell populations (Supplementary Table S4 and S5, Supplementary Figure S3).

**Figure 3.**
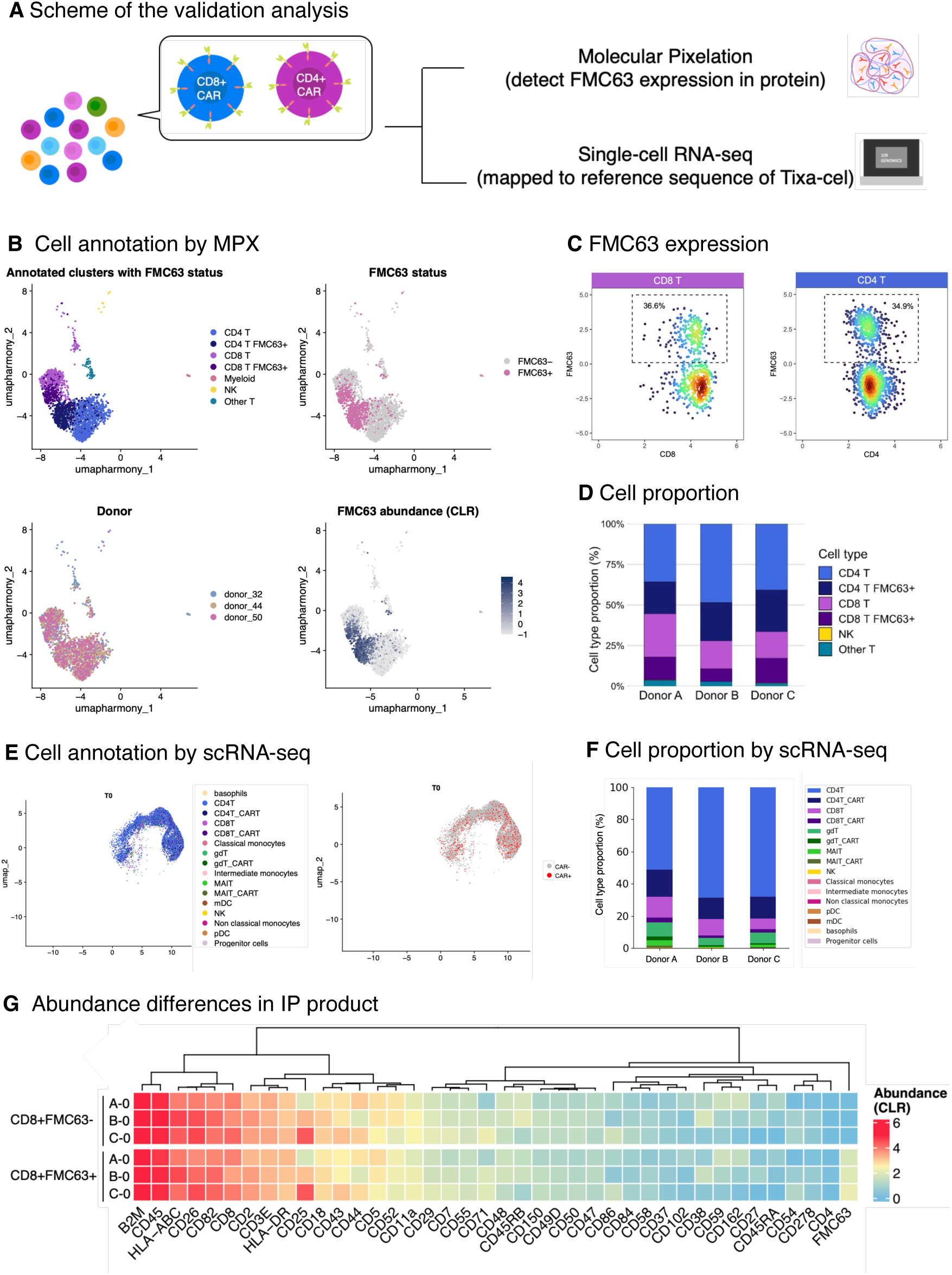
Characterization of clinical samples by protein abundance and single-cell transcriptome. **a,** Schematic overview of the analysis strategy using MPX and scRNA-seq. **b,** Cell type annotation of clinical samples based on MPX protein profiles. **c,** FMC63 abundance in CD4^+^ and CD8^+^ T cell populations measured with MPX. **d,** Cellular composition determined by MPX analysis. **e,** Identification of CAR-T cells using single-cell RNA-seq. **f,** Cellular composition determined using single-cell RNA-seq. **g,** Heatmap showing differences in surface protein abundances of FMC63^-^CD8^+^ T cells (top) and FMC63^+^CD8^+^ T cells (bottom) in pre-infusion CAR products. All datasets shown in Figure 3 were derived from CAR pre-infusion products (A-0, B-0, and C-0).

Focusing on pre-infusion samples in MPX datasets, after excluding platelets, remaining cells were composed of 34.5 % CD8 T cells, 41.6 % CD4 T cells, 2.7 % other T cells, 16.7 % myeloid, and 4.5 % NK cells (Figure 3B). FMC63^+^ cells were defined as cells with an FMC63 ≥ 0 CLR expression score per cell (Figure 3C). In the pre-infusion CAR-T products, FMC63^+^ cells accounted for approximately 36 % of CD8^+^ and 35 % of CD4^+^ T cells, with cellular composition comparable across donors (Figure 3D).

We performed a similar characterization analysis using the scRNA-seq data. FMC63^+^ T cells were identified by mapping sequencing reads to the tisa-cel reference sequence and were detected in both CD8^+^ and CD4^+^ T cell subsets (Figure 3E). The proportions of FMC63^+^ cells were consistent with those in MPX results (Figure 3D and 3F). CAR^+^ cells were detected in 18.1 % of CD8^+^ and 20.3 % of CD4^+^ T cells by scRNA-seq across pre-infusion samples. Notably, the MPX data showed a higher frequency of FMC63^+^ T cells, suggesting a higher sensitivity for detecting CAR^+^ T cells than that of single-cell RNA-seq (Figure 3D and 3F), which is consistent with a previous report that conventional single-cell RNA-seq shows limited sensitivity for CAR detection^19^. This discrepancy may also reflect differences between protein and mRNA expression levels.

The surface proteins on FMC63^+^CD8^+^ T cells were examined. As shown in a heatmap (Figure 3G), protein expression profiles were largely concordant across patients, suggesting robust detection with this method, as well as the stable production of CAR-T products. We also confirmed the abundance of these proteins in FMC63^-^CD8^+^T cells (Figure 3G). These data demonstrate that protein abundances in FMC63^+^ and FMC63^-^ cells was almost the same. Of note, CD25 showed relatively high expression in both FMC63^+^ and FMC63^-^ T cells from donor C.

### Surface polarity and spatial distribution in pre-infusion CAR-T cells

Leveraging unique capabilities of the MPX assay, the distribution of FMC63 and other surface proteins on FMC63^+^CD8^+^ T cells was analyzed. For each marker in each cell, a polarity score, which is a measure of biased spatial location, was calculated based on Moran’s *I* autocorrelation statistics using a cell adjacency graph. A positive score indicates a clustered distribution, whereas scores near zero reflect a random pattern^20^.

A heatmap of the polarity scores showed a largely comparable surface distribution pattern across pre-infusion samples from the three donors (Figure 4A). Consistent with immunofluorescence studies conducted prior to infusion^21^ and our in vitro observations, CAR molecules did not exhibit significant polarization prior to infusion (Figure 4B). In contrast, CD82, for which cluster formation is reported to occur upon T cell activation^22^, displayed consistently high polarity scores across all donors, indicating the highly active cytotoxic state of these products (Figure 4B).

**Figure 4.**
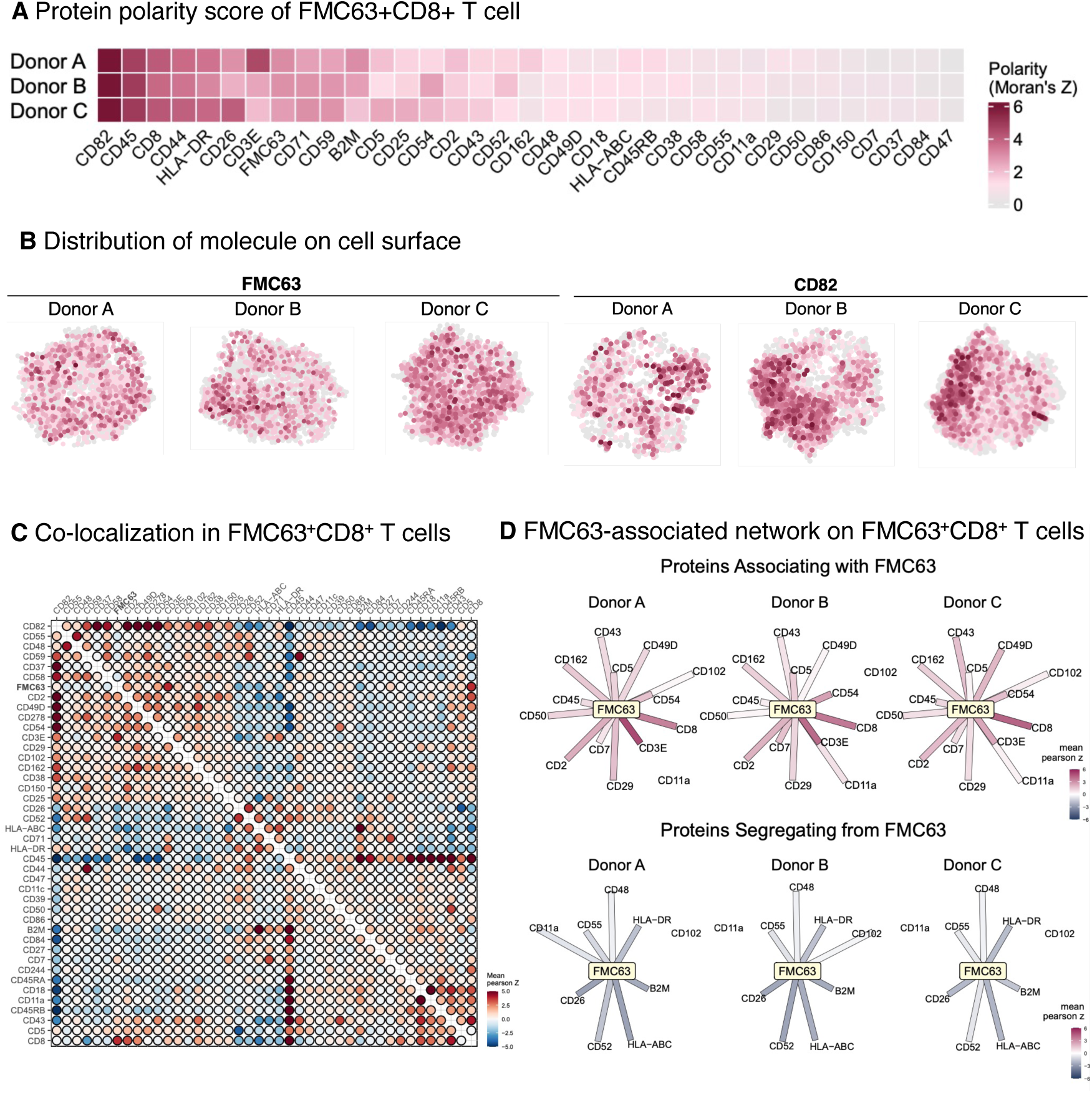
Characterization of CAR-T products by polarity and colocalization. **a**, Average polarization scores of selected markers on FMC63^+^CD8^+^ cells from three donors. **b,** Representative MPX images showing uniform distribution of FMC63 and polarized localization of CD82 **c,** Heatmap of average protein co-localization profiles of FMC63^+^CD8^+^ cells across the three donors. **d,** Network representation of proteins showing positive and negative co-localization with FMC63.

A key advantage of the MPX assay is its ability to assess protein co-localization on cell surfaces. A co-localization score was calculated for each protein, and these scores were compared across different donors to evaluate averaged co-localization (Figure 4C). Several known co-receptors (e.g., CD3, CD8, CD2, and CD5) and adhesion proteins (e.g., CD49d and CD54 (ICAM-1)) exhibited strong co-localization patterns (Figure 4C), supporting the technical robustness of spatial analysis using MPX. Further, inter-donor variability was minimal, indicating high reproducibility.

In the CAR-centered protein interaction network (Figure 4D), FMC63 consistently colocalized with CD8, CD3E, and other proteins. This network, encompassing multiple TCR-associated components, appeared to be highly similar across donors and was partly similar to that observed in an orthogonal preclinical CAR-T cell model.

### Longitudinal changes in cellular composition and protein abundance following CAR-T infusion

We next examined longitudinal changes following infusion. FMC63^+^ T cells were present in both CD8^+^ and CD4^+^ T cells in the pre-infusion samples (A-0, B-0, and C-0). Following infusion (A-1, A-2, A-3, B-1, and C-1), NK cell and myeloid cell populations increased, reflecting immune system reconstruction in the donors (Figure 5A). In addition, populations of FMC63^+^ cells changed dynamically over time. In donor A, FMC63^+^CD8^+^ T cells increased from 33.8 % (A-0) to 57.0 % (A-1), whereas FMC63^+^CD4^+^ T cells decreased from 35.3 % to 16.0 %. The number of FMC63^+^ T cells in this donor peaked on day 8 and then gradually declined (Figure 5B). Donor B showed a relatively stable FMC63^+^CD8^+^ cell population (31.3 % to 29.1 %) but a decrease in the FMC63^+^CD4^+^ cell population (33.1% to 11%). Donor C exhibited decreases in both FMC63^+^CD8^+^ (48.7 % to 32.0 %) and FMC63^+^CD4^+^ (39.0 % to 7.9 %) fractions (Figure 5B). scRNA-seq data showed similar trends, although the MPX assay showed higher sensitivity and detected a greater fraction of CAR^+^ T cells (Figure 5C and Supplementary Figure S4). PhenoCycler-Fusion quantification confirmed CAR^+^ CD8^+^ T cell expansion on day 8 in donor A (Supplementary Figure S5).

**Figure 5.**
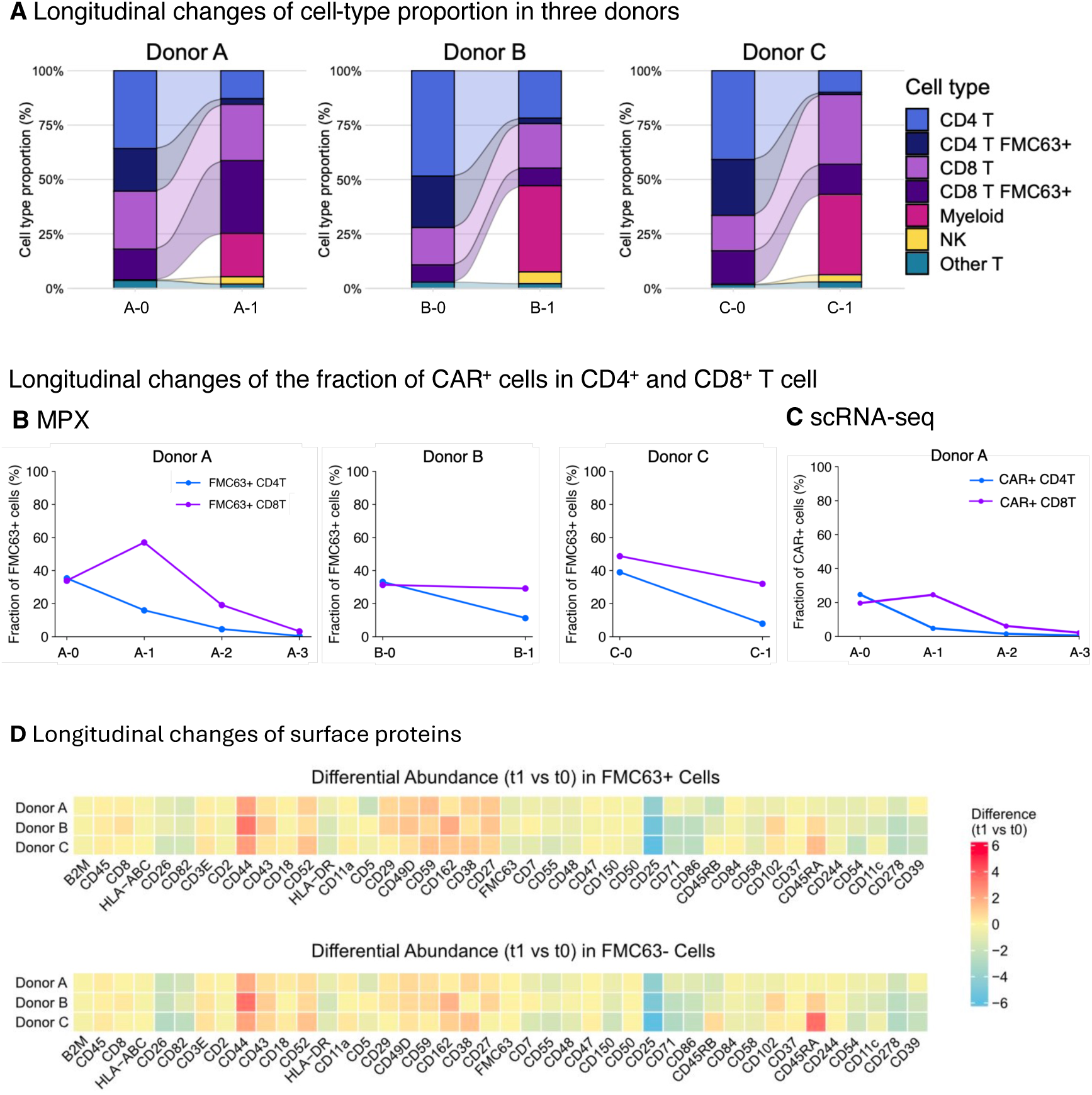
In vivo-induced changes in cellular composition and protein abundance. **a,** Longitudinal changes in cellular composition across the three donors following CAR^+^ T cell infusion. **b,** Longitudinal changes in the proportion of CAR^+^ T cells from donors A, B, and C measured with MPX. **c,** Corresponding longitudinal dynamics measured by single-cell RNA-seq of samples from donor A**. d,** Heatmap showing longitudinal changes in surface protein abundances of CD8^+^ CAR^+^ T cells across donors.

Despite these dynamic changes, surface protein abundance in pre- and post-infusion samples was largely similar across donors (Figure 5D). Although several changes were observed, no clear features distinguished the patients with sustained responses (donors A and B) from those with relapse (donor C). Similarly, the FMC63 polarity analysis did not reveal notable longitudinal differences between donors (Supplementary Figure S6), suggesting that clinical differences may not be fully explained by changes in protein abundance.

### Longitudinal remodeling of CAR-centered molecular co-localization networks

Longitudinal changes in FMC63-associated molecular organization were examined, in addition to changes in protein abundance. Donor B was excluded from this analysis due to insufficient post-infusion FMC63^+^CD8^+^ T cells (Figure 6A).

**Figure 6.**
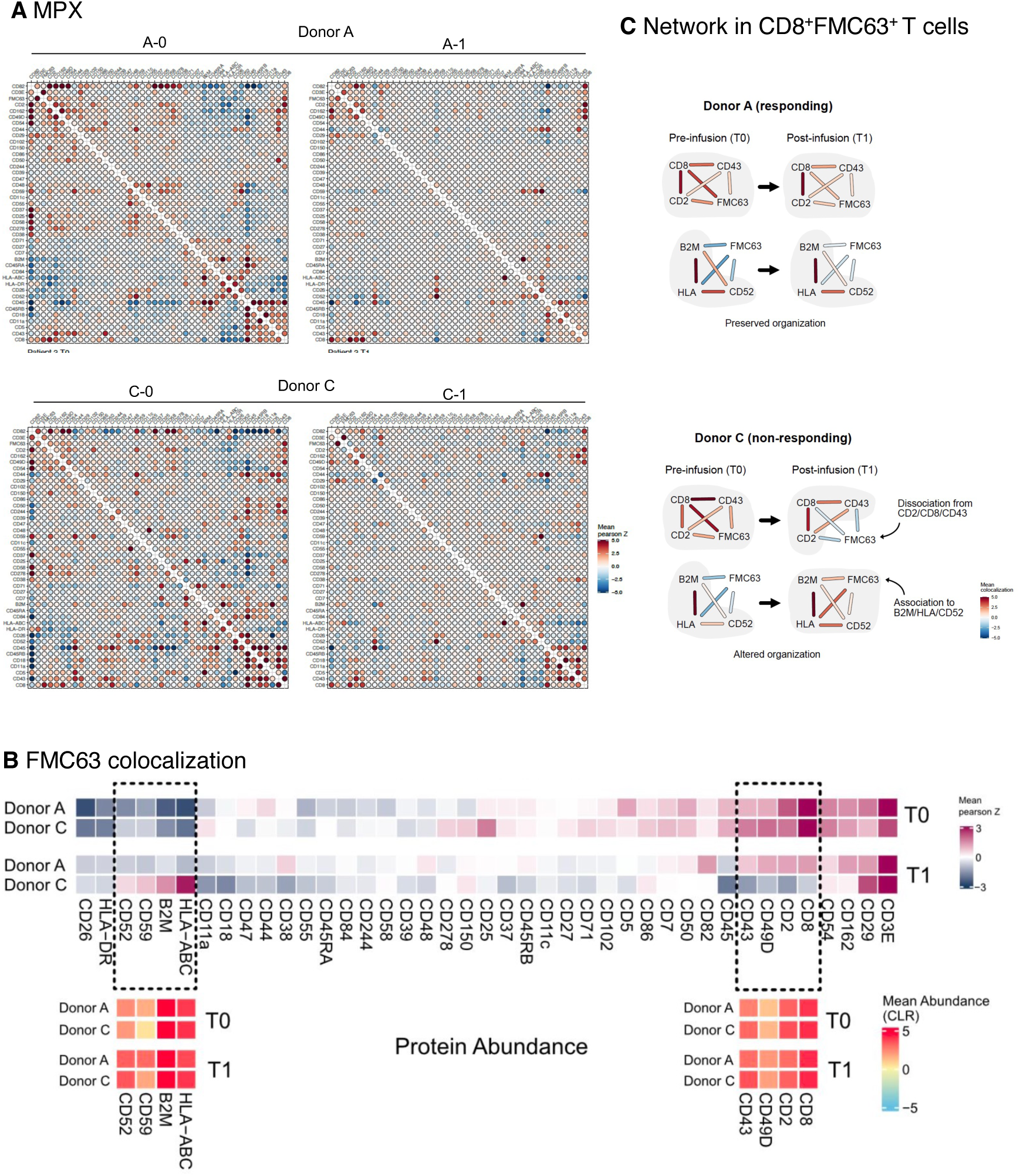
In vivo-induced remodeling of CAR-centered protein organization in CD8^+^ FMC63^+^ cells. **a,** Average protein co-localization heatmaps for donor A (top) and donor C (bottom) in the pre-infusion CAR-T product (A-0 and C-0) and post-infusion PBMC samples (A-1 and C-1). Several co-localization changes are shared between donors, including relocation of CD82, characterized by reduced co-localization with CD2 and CD3e and increased colocalization with CD8. **b,** Changes in FMC63-centered protein co-localization scores between pre-infusion CAR-T products (A-0 and C-0) and post-infusion PBMC samples (A-1 and C-1). **c.** FMC63-associated networks in donors A and C. Pre-infusion FMC63 co-localized with CD8, CD2, and CD43 in both donors. Post-infusion, donor C showed reorganization of FMC63 interactions and formation of a novel network with HLA, B2M, and CD52, whereas donor A maintained the original network.

The average co-localization network heatmap indicated many post-infusion changes were shared between donors A and C, reflecting conserved reorganization of immune protein architecture following CAR-T cell infusion. For example, co-localization of FMC63 with CD2 and CD3E was markedly reduced, whereas its co-localization with CD8 was enhanced with CD82 (Figure 6A).

However, donor- and time-specific differences were observed in the FMC63-centered network (Figure 6B and 6C). In donor A, network FMC63-CD2-CD8-CD43 from a pre-infusion (A-0) sample was retained after infusion (A-1). Donor C also exhibited this network prior to infusion (C-0). However, FMC63 dropped from this network after infusion (C-1), suggesting that active CAR-T cell function was lost at this stage. Instead, a novel CAR-CD52-B2M-HLA network was formed (C-1). This network, seemingly consisting of proteins unrelated to cytotoxicity, was absent from donor A samples (A-0 and A-1) and the pre-infusion donor C sample (C-0) (Figure 6C). Importantly, no significant differences were observed in the expression levels of these proteins between donors A and C pre- and post-infusion (Figure 6C). Immunosuppressive interventions, including the use of corticosteroids in addition to tocilizumab, may have further influenced the post-infusion remodeling of CAR-associated membrane organization, potentially promoting dissociation of FMC63 from the CD8-CD2 functional network and contributing to decoupling of inflammatory responses from sustained antitumor activity. High-dose and prolonged corticosteroid exposure have both been reported to be associated with inferior clinical outcomes after CAR-T cell therapy, likely due to impacts of this exposure on CAR-T cells and the surrounding immune microenvironment^23^. Notably, our findings suggest that even relatively limited corticosteroid use, as observed in this cohort, may induce qualitative changes in CAR-T cell functional architecture that are detectable at the membrane protein organization level.

Collectively, these results suggest that MPX including FMC63 AOC enables the detection of CAR-centered protein interaction networks that are not captured by conventional protein abundance measurements or scRNA-seq. Such spatial reorganization may underlie differential therapeutic responses and highlight the unique capability of MPX to resolve protein architecture at single-cell resolution.

## Conclusions

In this study, we developed and validated an assay to monitor the molecular dynamics of CAR-T cells. An FMC63-targeting AOC was generated and integrated into a 76-plex AOC panel. This framework enabled simultaneous quantification of protein abundance, spatial organization of CAR molecules, and co-localization of these molecules with other surface proteins at the single-cell level. By applying this framework to longitudinal clinical specimens, we showed that remodeling of CAR-centered co-localization networks on the surface of CD8^+^CAR^+^ T cells, rather than differences in CAR protein abundance, was a key difference between effectively treated and relapse cases.

This study had several limitations. First, the cohort size was limited to three patients, precluding statistically powered conclusions and generalization of clinical associations. Therefore, the reported networks should be interpreted as hypothesis-generating observations rather than definitive biomarkers of therapeutic outcomes. Second, although co-localization networks were quantified at single-cell resolution, functional validation of specific molecular interactions was not performed. To the best of our knowledge, no existing analytical method enables equivalent spatial network analyses at this scale. Third, independent replication in external cohorts will be required to assess the reproducibility and predictive value of the demonstrated approach. Despite these limitations, our study establishes a technical framework for systemic characterization of surface protein organization in CAR^+^ T cells under clinical conditions. By integrating protein abundance and spatial assays, MPX expands the analytical resolution of single-cell profiling beyond that of conventional methods and provides a complementary layer of information for studying cellular therapies.

Notably, protein abundance and polarity profiles were highly consistent across donors and time points, indicating the robustness of the assay. In contrast, remodeling of CAR-associated co-localization and networks distinguished relapse from non-relapse cases, suggesting that molecular organization may have contributed to clinical responses.

Future studies in a larger patient cohort, together with functional validation analyses using perturbation experiments, will be required to determine whether CAR-centered network organization represents a generalizable determinant of therapeutic efficacy and resistance. Taken together, these findings suggest that spatial organization of CAR-associated proteins, rather than protein abundance or polarity alone, represent an important determinant of CAR-T cell therapeutic responses, highlighting the value of MPX for dissecting molecular architectures through the course of cellular therapies.

## Methods

### Human subject research

Clinical samples were obtained from patients at Tsukuba University Hospital, Japan. This study was approved by the Ethics Committee of Tsukuba University Hospital (Project No. H24-075) and conducted in accordance with the Declaration of Helsinki.

### Data availability

The raw data have been submitted to the National Bioscience Database Center, Japan (J-DS001939-001).

### Flow cytometry analysis of CAR-T cells

CD19-targeting CAR-T cells (ProMab, Cat# PM-CAR1007) and peripheral blood mononuclear cells (PBMCs) from a healthy donor were stained with 0.04–5 μg/mL anti-FMC63 antibody probe (Pixelgen Technologies) for 40 min at 4 C°. Cells were washed and then were incubated with 2 μg/ml phycoerythrin (PE)-conjugated secondary antibody for 40 min at 4 C°. Cells were processed on a CytoFLEX (Beckman Coulter) and analyzed using FlowJo software.

### Co-culture of Raji cells and CD19-targeting T cells

Raji cells (DSMZ, Cat# ACC 603) and CD19-targeting CAR-T cells were used in the co-culture assay. Cells were maintained in RPMI medium (ThermoFisher, Cat# 61870-010) supplemented with 10 % fetal bovine serum (ThermoFisher, Cat# A5670801) and 1 % penicillin-streptomycin (Gibco, Cat# 15140148). Cells were co-cultured at an effector-to-target (E:T) ratio of 5:1, and samples were collected 4 or 24 h after the initiation of co-culture. Following collection, the cells were cryopreserved and stored until subsequent molecular pixelation analysis.

### Multiplexed immunostaining by PhenoCycler-Fusion

To validate the FMC63 antibody for detection of CAR-T cells, PBMCs from a healthy donor with no history of CAR-T cell therapy (negative control) and donor A (A-0; CAR-T cell-positive, positive control) were analyzed. The cells were stained using a PhenoCycler-Fusion Sample Kit (Cat#SKU7000017, Quanterix). FMC63 carrier-free monoclonal anti-FMC63 antibody (Cat# FM3-Y45A1, AcroBiosystems) was conjugated to the BX134 barcode (Cat#SKU5550030, Quanterix) using an Antibody Conjugation Kit (Cat#SKU7000009, Quanterix). Together with other antibodies, a panel of 19 antibodies targeting PBMC subsets was designed (antibody lists and targets are summarized in Supplementary Table S2).

Cells were fixed with Hydration Buffer (Quanterix) containing 1.6 % paraformaldehyde at room temperature for 20 min, washed with additional Hydration Buffer, and smeared onto glass slides. After air-drying, the glass slides were washed twice with PBS (Cat# 045-29795, FUJIFILM) and equilibrated in Staining Buffer (Quanterix) for 20 min at room temperature. Glass slides were stained at room temperature for 3 h using the antibody panel. After staining, the samples were processed according to the fresh-frozen staining protocol and imaging was performed using a PhenoCycler-Fusion system following the manufacturer’s instructions (PhenoCycler-Fusion User Guide_2.1.0, PD-000011 Rev L). The output data were visualized using QuPath^24^ (v0.6.0).

### Quantification analysis of PhenoCycler-Fusion data

For the quantitative analysis, fluorescence intensity values for each channel were extracted from qptiff files at the single-cell level following cell segmentation in QuPath using StarDist^25^. These values were log10 transformed prior to downstream analysis. The threshold for each marker was determined by inspecting the intensity distribution. Thresholds were fixed for each marker and applied uniformly across all samples included in the analysis to ensure reproducible and comparable classification of positive and negative cells. The threshold for FMC63 was further validated using a sample from a negative control, a heathy donor, in which fewer than 0.2 % of cells were classified as positive (Figure S2), thus supporting the specificity of the selected threshold.

Based on these thresholds, the cells were classified as positive or negative for each marker, and the number and proportion of positive cells were calculated for each sample. To identify the CAR-expressing T cell populations (Figure S3), cells exhibiting triple-positive signals (FMC63^+^CD3e^+^CD8^+^ and FMC63^+^CD3e^+^CD4^+^) were defined as CAR^+^CD8^+^ and CAR^+^CD4^+^ T cells, respectively, using a predefined threshold.

### Molecular pixelation library preparation and sequencing

Cells were processed according to the manufacturer’s protocol (v2 protocol PXGIMM002, Immunology panel 2 Human v2, Cat# PXGIMM002PP, and DP00002 CAR Antibody-Oligo Conjugate spike-in (v1.00)). Constructed libraries were sequenced using either an Illumina NovaSeq6000 or Element AVITI system (Read1: 44 cycles; i7: 8 cycles; i5: 8 cycles; Read2: 78 cycles) with 15 % PhiX spike-in. Sequencing statistics and additional details are summarized in Supplementary Table S4.

### Initial data processing of MPX after sequencing

Sequencing data were initially processed using nf-core/pixelator (v1.4.0)^20,26^. The pipeline was executed with NextFlow and Singularity using their default parameters. A custom panel file (human-sc-immunology-spatial-proteomics-2-cart), which included an FMC63-targeting antibody oligonucleotide conjugate, was used as input.

### MPX data analysis

MPX-generated sequencing data were initially processed using nf-core/pixelator (v1.4.0)^20,26^. The pipeline was executed with NextFlow and Singularity using their default parameters. A custom panel file (human-sc-immunology-spatial-proteomics-2-cart), which included an FMC63-targeting antibody oligonucleotide conjugate, was used as input. Briefly, raw sequencing reads were subjected to quality control and filtering prior to assembly into amplicons that captured protein interactions. Amplicons were then grouped by marker assignment and error-corrected to produce an edge list representing a full sample as a molecular graph. Individual cells were resolved from this graph using Leiden community detection, and only the components representing intact cells were retained for downstream analyses. Spatial metrics were subsequently computed for each cellular component and stored together with the associated data in PXL format.

Downstream analyses were carried out in R v4.4.3 using pixelatorR v0.14.0 (https://github.com/PixelgenTechnologies/pixelatorR), pixelatorES v0.6.0 (https://github.com/PixelgenTechnologies/pixelatores), and Seurat v5.3.0. Data visualization was performed using ggplot2 v3.5.2 and ComplexHeatmap v2.22.0.

Components were filtered based on multiple quality metrics: (1) count thresholds requiring 10,000–100,000 unique molecular identifiers (UMIs) per component, (2) normal Tau [2] type classification to exclude potential antibody aggregates, (3) isotype control antibody fraction (mIgG1, mIgG2a, mIgG2b) <1% of total molecules, and (4) ACTB fraction <1% to exclude permeabilized or damaged cells. Protein abundance data were normalized using a centered log-ratio (CLR) transformation and then scaled. Principal component analysis (PCA) was performed, and the top 15 principal components were selected based on explained variance. Batch effects across samples were corrected using Harmony integration of sample identities. Uniform Manifold Approximation and Projection (UMAP) was computed using Harmony-corrected dimensions for visualization. Cell clusters were identified using a shared nearest neighbor (SNN) graph constructed from Harmony (v1.2.3)-corrected dimensions, followed by Louvain clustering at a resolution of 0.5. Cluster marker proteins were identified using Wilcoxon rank-sum test. Clusters were manually annotated as major cell populations (CD4^+^ T cells, CD8^+^ T cells, NK cells, myeloid cells, platelets, and other T cells) based on canonical marker expression. FMC63-positive cells (CAR-T cells) were identified using a CLR-normalized abundance threshold ≥0.5, and cell type annotations were combined with FMC63 status for downstream analysis.

Differential protein abundance, polarization, and colocalization were assessed using pixelatorR functions RunDAA(), RunDPA(), and RunDCA(), respectively. For abundance, CLR-normalized values were used. Protein polarization was quantified using Moran’s Z-scores, with data filtered to retain proteins with >50 counts per cell and detected in >20 cells per condition. For the colocalization analysis, Pearson Z scores, filtered to include marker pairs where the lower-abundance marker exceeded 50 counts per cell, were applied, and pairs were detected in ≥50 cells per sample.

### Identification of B cell patches on co-cultured CAR-T cells

We implemented a graph-based patch-detection algorithm to identify discrete B cell patches. First, node-level count matrices were expanded by aggregating expression values from immediate neighbors using an adjacency matrix of the cell graph. This step effectively smoothed local signals to better capture regional marker density.

Specifically, B cell identity was defined as the combined expression of CD20 and CD40. In the component graphs where these markers were detected, we initialized a label propagation algorithm where nodes were pre-labeled based on their aggregated marker counts. Nodes with a combined UMI count greater than 10 were assigned to one group, while all others were assigned to a second group.

Following label propagation, the least frequent community was identified as the candidate patch population, under the assumption that B cell aggregates represent a small subset of aggregates in the overall graph. To ensure robustness of the identified structures, candidate patches were filtered to include only connected components consisting of at least 40 nodes. Finally, patch sizes and relative proportions were calculated for each component to quantify the extent of B cell patching.

### Single-cell RNA-seq library preparation and sequencing

For single-cell RNA-seq, we employed 10× Genomics Single-cell Chromium 5 ′ platform (Next GEM Single Cell 5’ Kit v2). Libraries were prepared according to the manufacturer’s instructions (CG000331). Briefly, cells were diluted and loaded to target 10,000 cells per sample. After GEM generation, cDNA was amplified for 13 PCR cycles. For indexing, PCR was performed for 12 or 13 cycles depending on the library concentration. Sequencing was then conducted using an Element AVITI sequencing system (Read1:28 cycles; i7:10 cycles; i5:10 cycles; Read2:90 cycles). Sequencing statistics and further details are summarized in Supplementary Table S5.

### Read mapping and CAR read identification after sequencing for scRNAseq

After sequencing, CellRanger (version 9.0.1) was used for initial processing. Reads were mapped to reference genome (GRCh38-2024-A & vdj_GRCh38_alts_ensembl-7.1.0). To identify CAR-T cells, unmapped reads (flag = 4) with valid cell barcodes (CB) and UMIs (UB) were extracted from CellRanger-generated BAM. These reads were mapped to a customized Kymriah reference sequence derived from a previous study^11^ using BWA-MEM^27^ (version 0.7.17-r1188, -M options). After mapping, reads that passed the filter (AS ≥ 80 & mapq > 50) were identified as CAR reads.

### Quality control and preprocessing

Downstream analysis was performed using R package Seurat^28^ (version 5.1.0). For quality control, cells were filtered using the following criteria: nFeature_RNA > 1000 & percent.mt < 30 & percent.rib > 2 & exon_rate > 35. Percent.mt (proportion of mitochondria genes) and percent.rib (proportion of ribosomal protein genes) were calculated using the Seurat function PercentageFeatureSet with pattern “^MT-” and pattern “^RP[LS]”, respectively. Exon rates were calculated from CellRanger BAM files by parsing “RE” tag and aggregating counts per cell based on CB/UMI pairs, with PCR duplicates removed using a custom Python script.

### Dimensionality reduction and clustering

Following quality control, each Seurat object was processed using the standard Seurat workflow. Data were normalized using NormalizeData, highly variable features were identifieid with FindVariableFeatures (selection.method = “vst”, nfeatures = 1000), and the data were scaled using ScaleData (features = all.genes). Linear dimensionally reduction was performed through PCA using RunPCA (npcs = 10). Cells were clustered using FindNeighbors (reduction = “pca”, dims = 1:10) followed by FindClusters (resolution = 0.5). Non-linear dimensionally reduction was conducted by applying UMAP with RunUMAP (reduction = “pca”, dims = 1:10).

### Cell type annotation and CAR-T identification

Cells were annotated to cell type using R package SingleR^29^ (version 2.4.1) with a reference dataset from celldex package^30^ (version 1.12.0); specifically, the PBMC dataset^31^ MonacoImmuneData was applied. The detailed groups were consolidated into representative cell types (Supplementary Table S6). After annotation, the cells were further stratified based on the presence of CAR-derived reads identified in the previous mapping step. Cells positive for CAR (CAR reads > 1) were labeled according to their T cell subtypes as CD4_CART, CD8_CART, gdT_CART (γδ CAR T cell), or MAIT_CART.

### Integration of longitudinal datasets

Longitudinal Seurat objects from the same individual were integrated. Seurat objects were merged and re-normalized using NormalizeData. Highly variable features were identified using FindVariableFeatures, and the data were scaled using ScaleData. PCA was performed in RunPCA (npcs = 10), followed by integration with Harmony using the Seurat function IntegrateLayers (method = HarmonyIntegration, orig.reduction = “pca”). Cells were clustered using FindNeighbors (reduction = “pca”, dims = 1:20) and FindClusters (resolution = 0.5), and UMAP visualization was performed using RunUMAP (reduction = “pca”, dims = 1:20).

## Acknowledgements

The authors thank all of the anonymous donors who provided samples for this study. This study was supported by JSPS KAKEN (JP25K18811) to Y.K. and AMED (JP23jf0126003) to Y.S.

## Authorship Contribution

Y. K. and H.O. wrote the manuscript and interpreted the data. K.M. and H.N. contributed to patient recruitment, follow-up, and study design. L.F. and S.P. analyzed the data. J.Z. and A.S. performed the experiments and analyzed the data. M.S.Y. and Y. S. supervised the study. All authors reviewed and approved the final version of the manuscript.

