## Supplementary Information for "Spatial profiling of CAR protein organization reveals in vivo remodeling during CAR-T therapy"

Supplementary Figures

This file includes

- Supplementary Figure S1 to S5
- Supplementary Tables S1 to S6

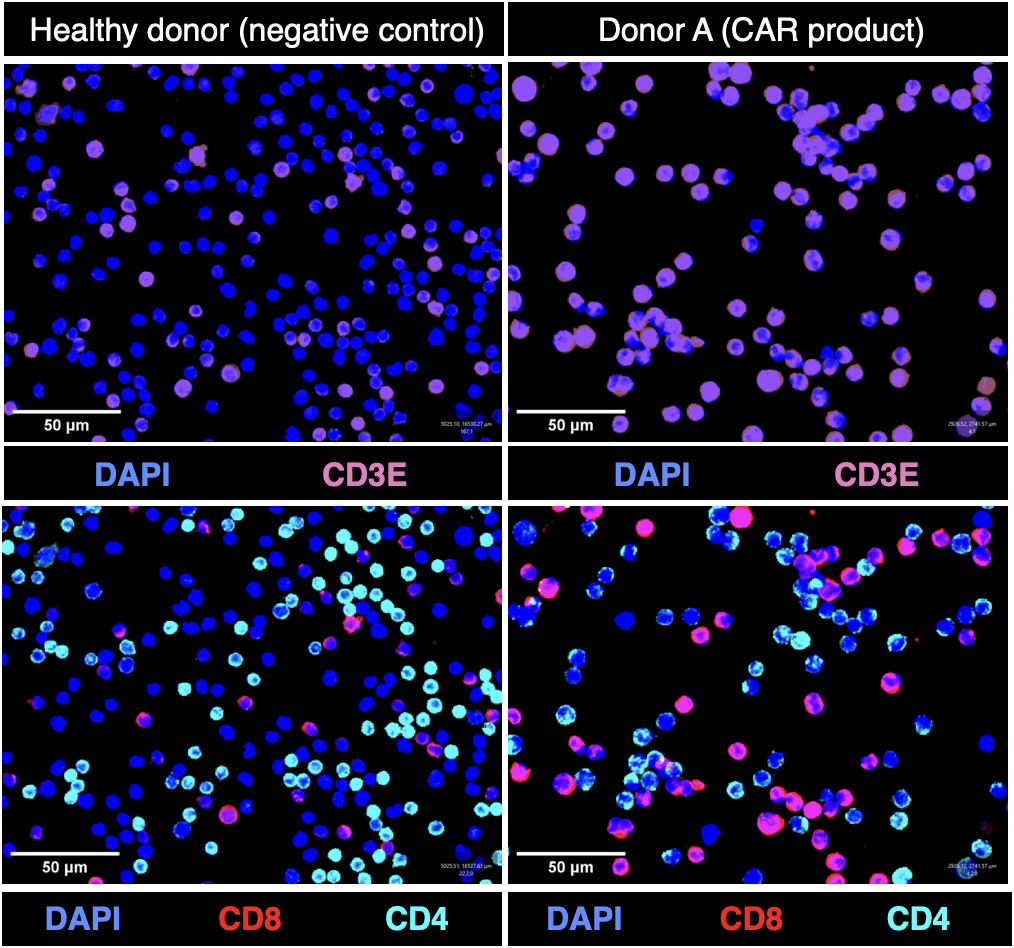

Fig. S1. PhenoCycler-Fusion images for selected representative T cell marker channels

Type or paste caption here. Create a page break and paste in the Figure above the caption.

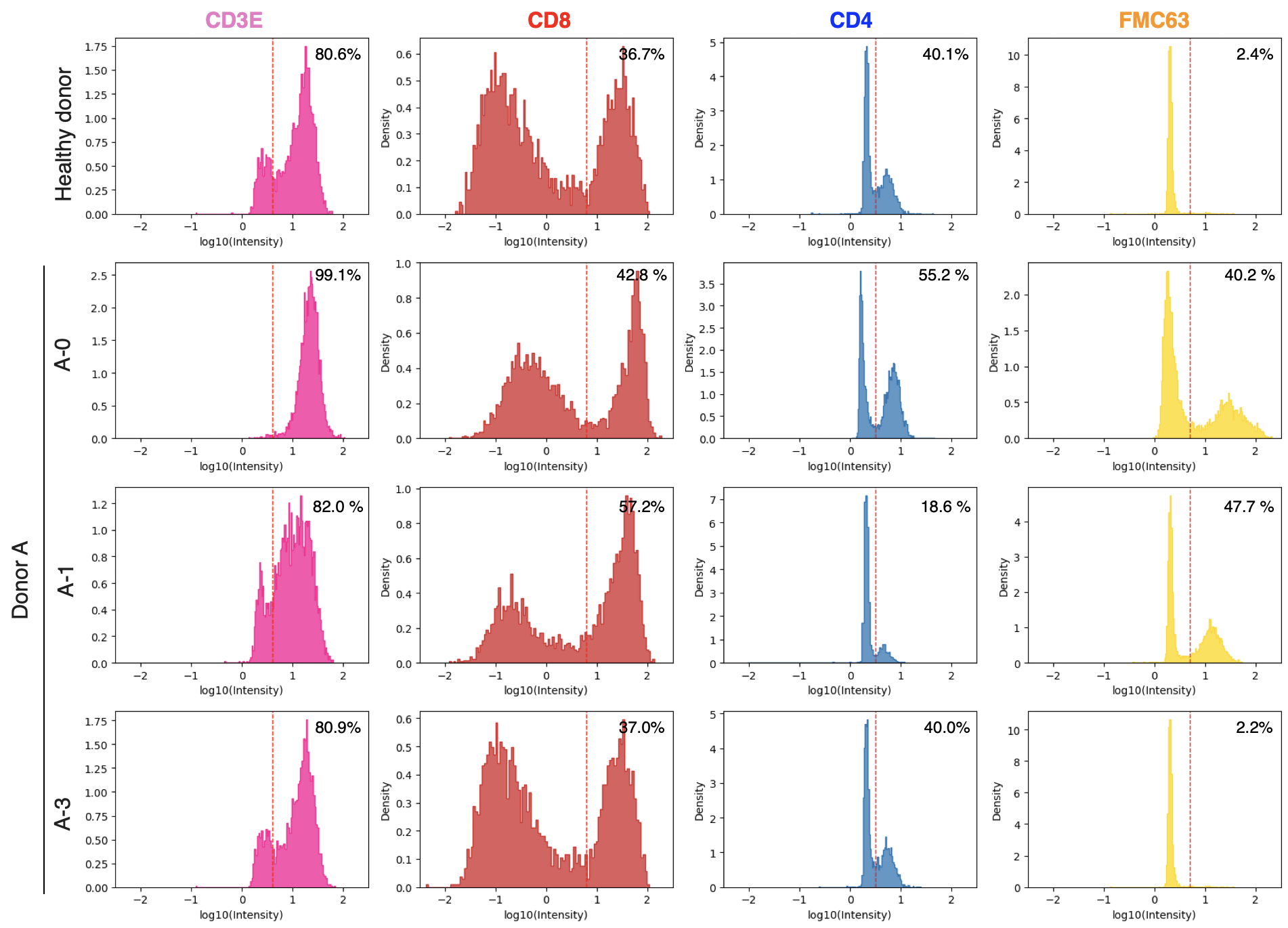

Fig. S2. Quantification of marker expression using PhenoCycler-Fusion

Distribution of log10-transformed intensity values for CD3E, CD8, CD4 and FMC63 across all samples, healthy donor, A-0, A-1 and A-3. Histogram shows single-cell intensity distributions for each marker. Dashed red lines indicates the threshold used to classify cells as positive and negative. Ratio of marker positive cells are shown on each plot.

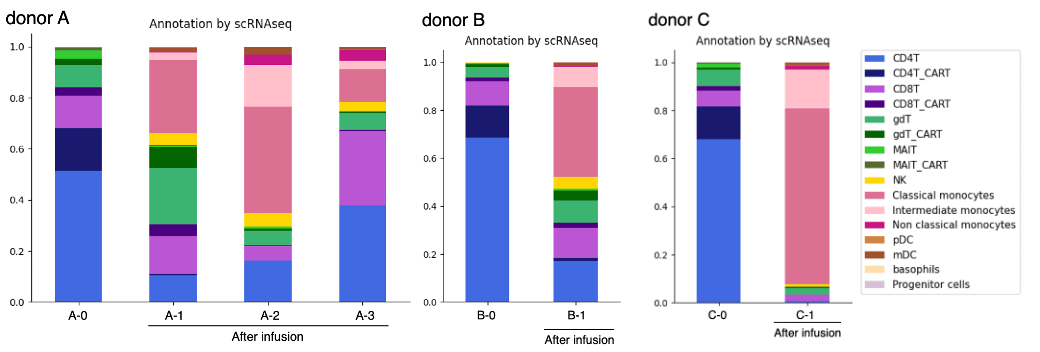

Fig. S3. Cell type proportions based on scRNA-seq

Longitudinal changes in cell type proportion in donor A, B and C are shown in bar plot. Cell types were annotated based on scRNA-seq data. Color indicates cell types, as shown in the accompanying color key.

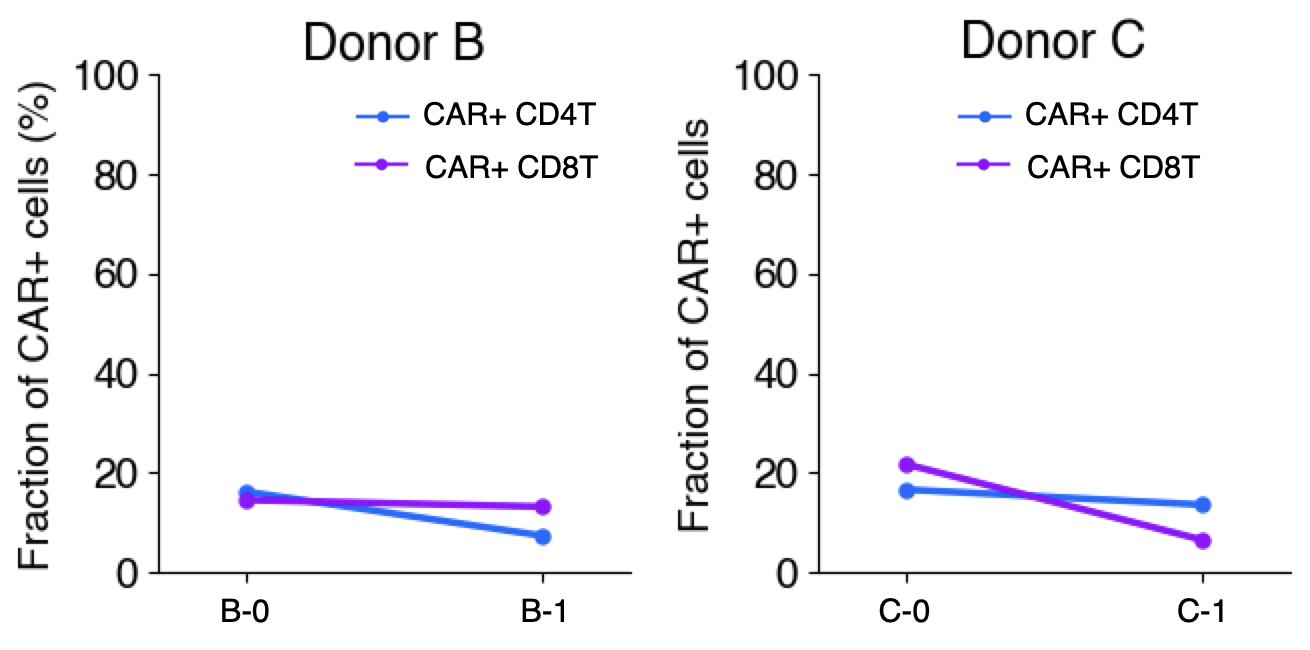

Fig. S4. Longitudinal changes of CAR^+^ CD4T and CD8T cells using scRNA-seq

Longitudinal changes of the percentage of CAR+ CD4T and CD8T cells in donor B and C are shown in line plot. Cell types were annotated based on scRNA-seq data. Color indicates cell types, as shown in the accompanying color key.

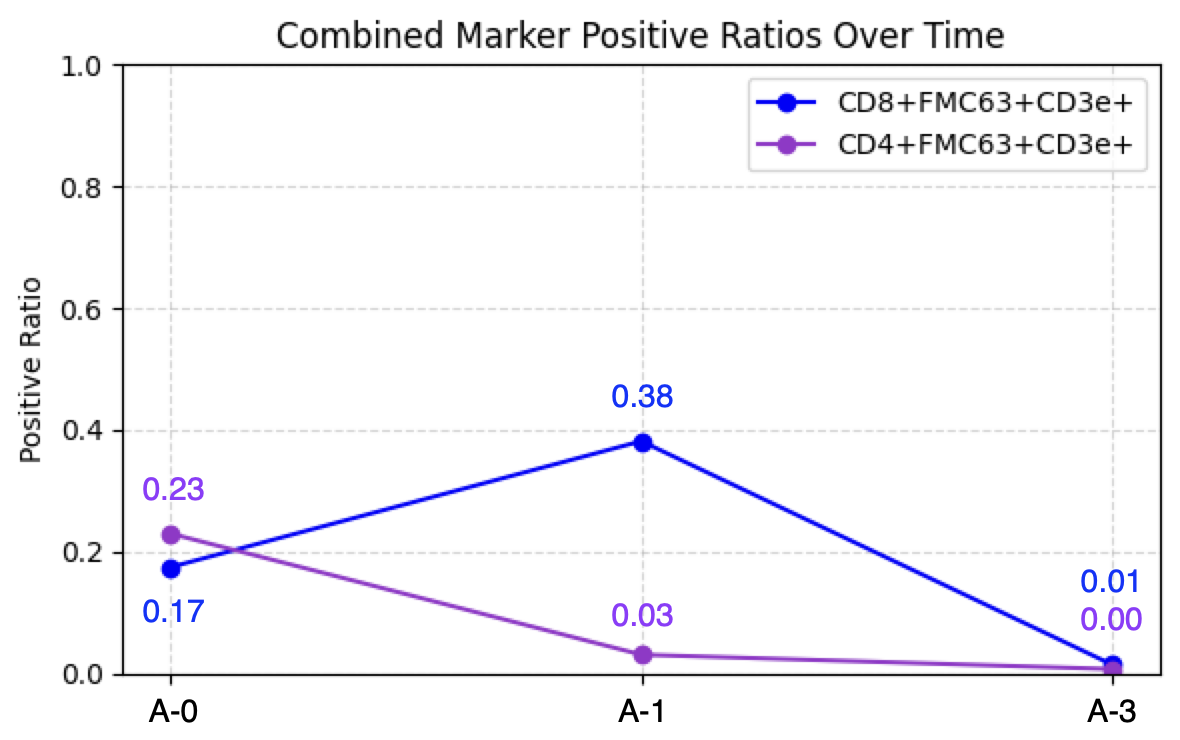

Fig. S5. Quantification of CAR-expression population using PhenoCycler-Fusion

Longitudinal changes in donor A. The proportion of triple-positive FMC63+CD3e+CD8+ cells (CAR+ CD8+ T cell) and FMC63+CD3e+CD4+ cells (CAR+ CD4+ T cell) were calculated for each sample and plotted across the time points (A-0, A-1 and A-3). The A-2 sample was not analyzed due to insufficient cell number. The ratio at each point is shown on the plot.

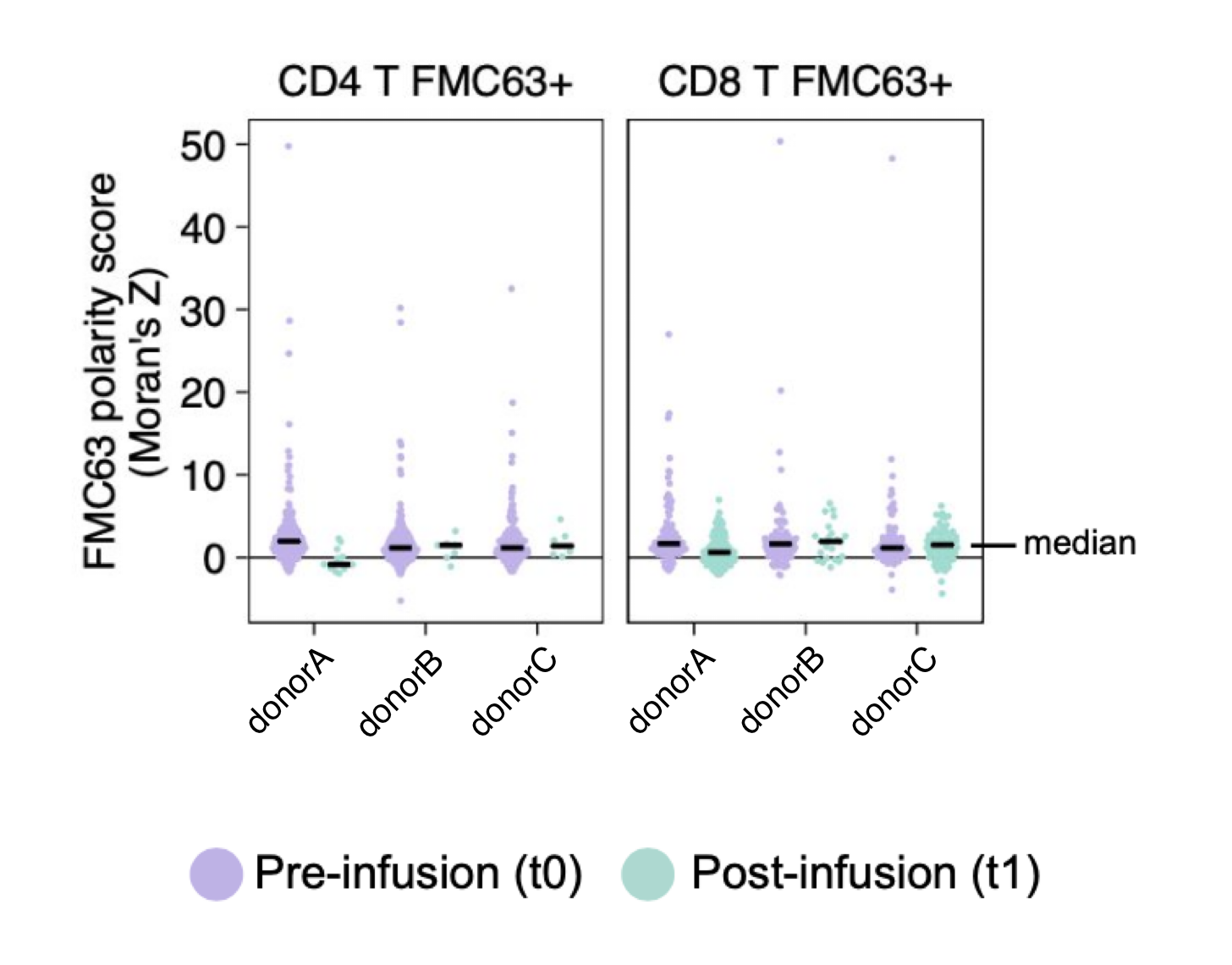

Fig. S6. Longitudinal changes of polarity score of FMC63 on CAR^+^ CD4T and CD8T cells

Longitudinal changes of the FMC63 polarity score (Moran’s Z) of CAR+ CD4T and CD8T cells in donor B and C are shown in bee swarm plot in three donors.

Table S1. Clinical information of the donors involved in the study

This table summarizes the clinical characteristics of the donor A, B, and C.

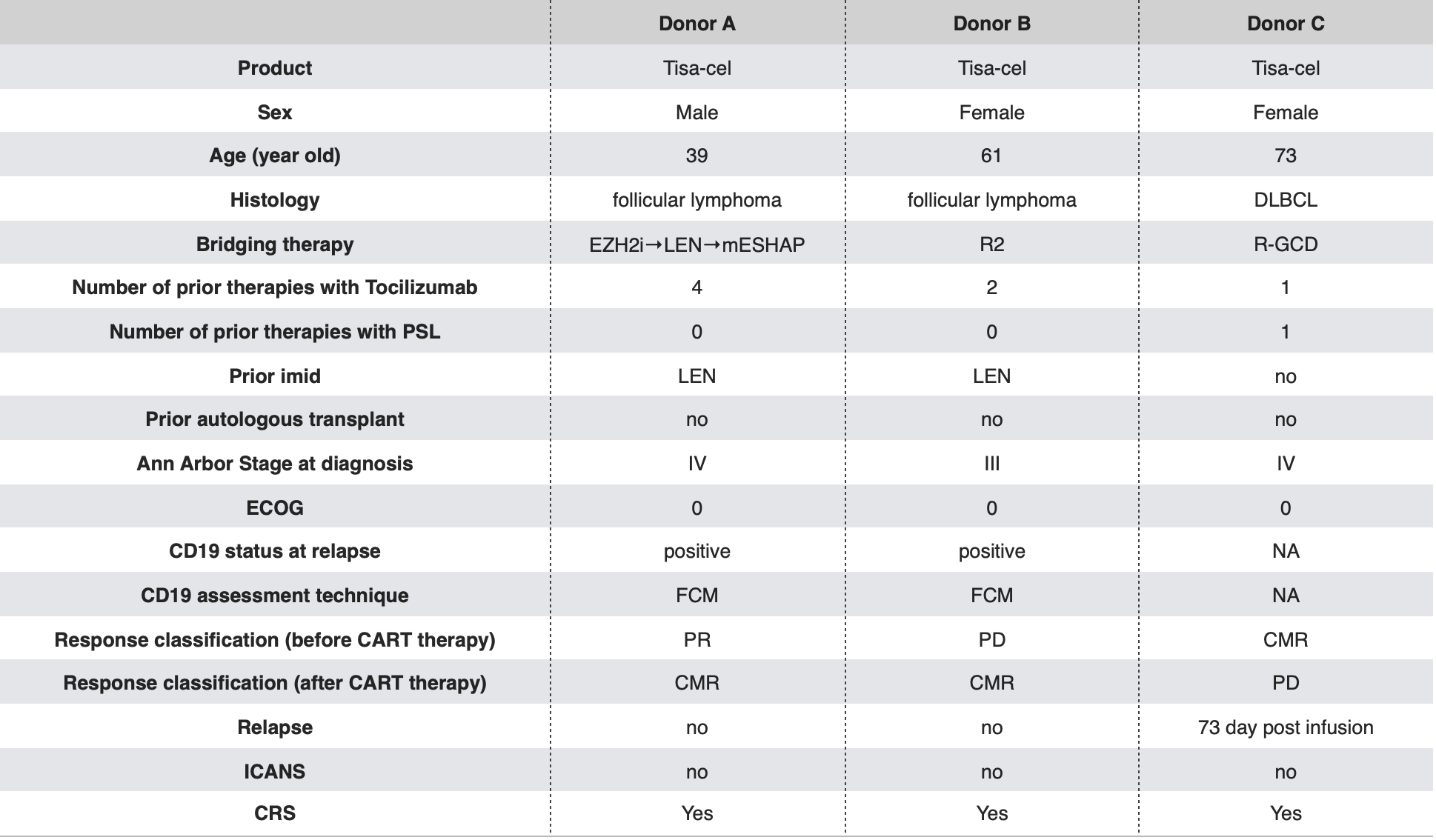

Table S2. List of antibodies in the panel for PhenoCycler-Fusion

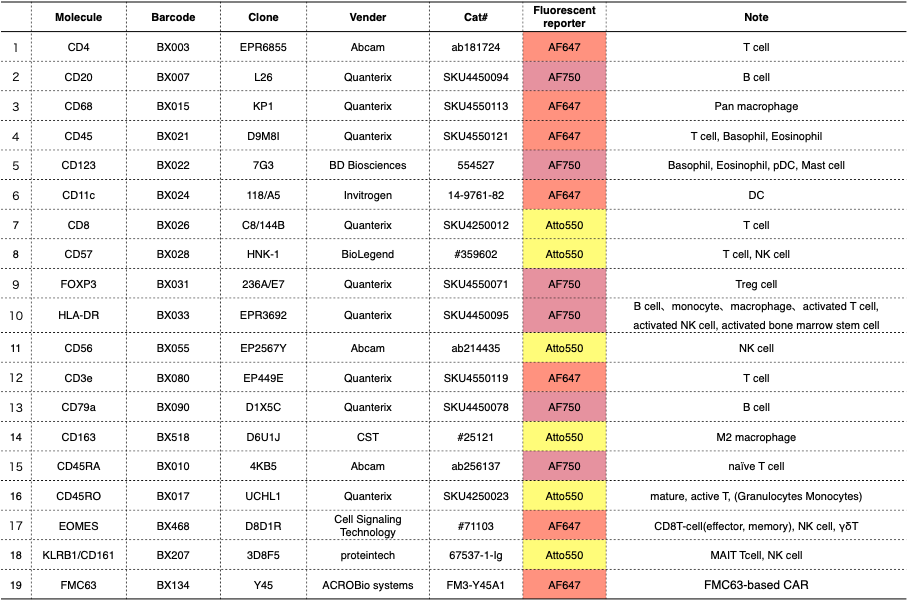

Table S3. Run design for PhenoCycler-Fusion

This table shows the run design for PhenoCycler-Fusion.

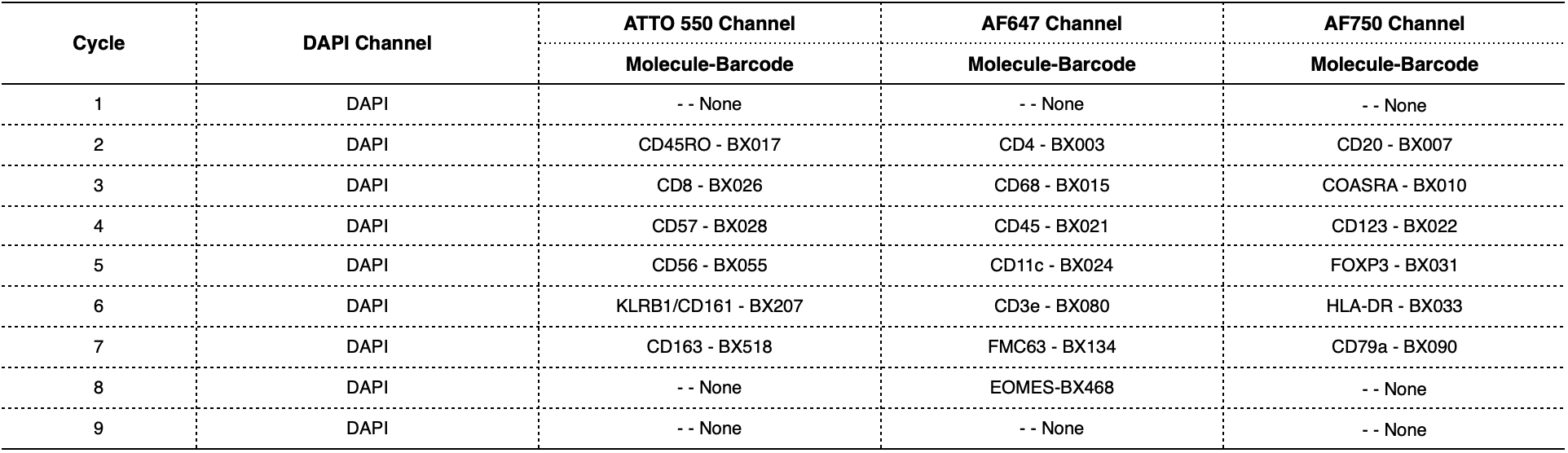

Table S4. Statistics for MPX libraries sequencing

This table summarizes the key sequencing metrics for the MPX libraries used in this study.

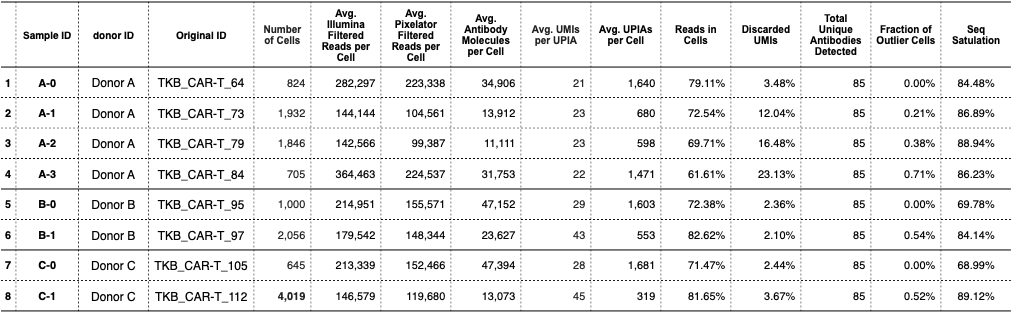

Table S5. Statistics of single-cell RNA-seq libraries sequencing

This table summarizes the key sequencing metrics for the scRNA-seq libraries used in this study.

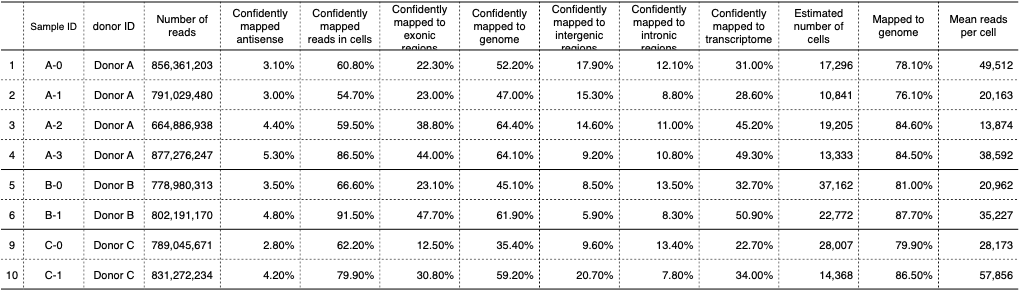

Table S6. Categories for cell type annotation

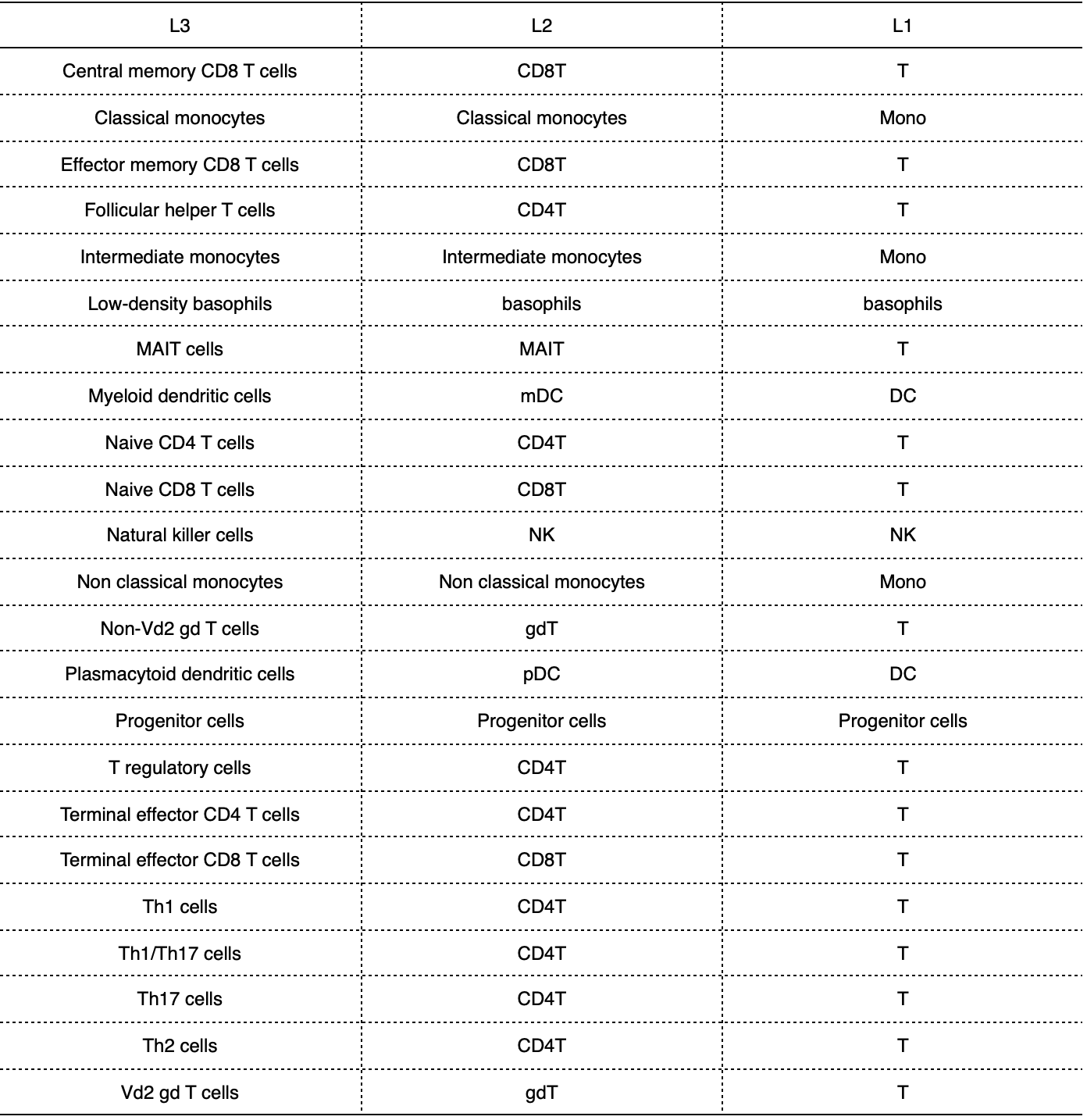
